# Virion encapsidation and cell attachment function of the reovirus attachment protein is influenced by its structural flexibility

**DOI:** 10.1101/2025.05.12.653539

**Authors:** Maximiliano L Garcia, Pranav Danthi

## Abstract

The reovirus σ1 attachment protein mediates virus interaction with host cell receptors that is critical for cell entry. Reovirus tropism is controlled by properties of σ1. σ1 is present as trimers that are held within turrets at the icosahedral vertices of reovirus virions. However, because σ1 has not been visualized on reovirus virions in high resolution structures and because the fulllength structure of purified σ1 protein has not been solved, it is not clear how σ1 is presented on virions. What properties of σ1 are essential for its incorporation on virions is also not known. In this study, we used ColabFold to model the structure of reovirus serotype 1 (T1) and serotype 3 (T3) σ1 proteins. We find that these proteins fold into similar structures with regions of flexibility between the head and body domains of σ1. We also predicted the structures of chimeric σ1 proteins comprised of domain swaps between T1 and T3 σ1 proteins. Our analyses indicate that chimeric proteins with mismatched body and head domain have increased flexibility in this region. Characterization of particles expressing such chimeric σ1 proteins demonstrated that deviation from the flexibility of parental σ1 leads to a reduction in σ1 incorporation on to the virion. Further, we find that even when incorporation is not affected, virus attachment to host cell receptors is influenced by altered σ1 flexibility. Finally, our work demonstrates that μ1 protein impacts the encapsidation pattern and receptor engagement properties of σ1 and that this effect is influenced by properties of the N-terminal portion of σ1.

**Importance:** Attachment to host cell receptors is a critical step in initiation of virus infection. Some viruses attach to cellular receptors via dedicated viral proteins. Both the number of attachment factors present on the virus and whether they are present on the virus particle in the correct form can influence cell attachment. Here, using reovirus as a model, we use a protein structure prediction algorithm to model the as yet unknown structure of full-length reovirus attachment protein σ1. We find predicted regions of flexibility in the protein and identify how this flexibility is regulated. We find that the flexibility of σ1 independently regulates whether it is stably incorporated into particles and if can efficiently interact with host receptors.

## Introduction

Since viruses are obligate intracellular parasites, attachment of virus particles to receptors on susceptible cells is a critical step in infection of all viruses (1). Viruses mediate this step via specific structural features on nonenveloped virus capsids. For example, canyon regions of the poliovirus capsid engage with the poliovirus receptor, CD155 (2). Alternatively, such an interaction occurs through the presence of dedicated proteins on the virion surface that are often referred to as attachment proteins. For example, the Spike protein of Coronavirus or fiber like structures found on adenovirus and mammalian orthoreovirus (3-5). In situations where the attachment protein is not integral to the formation of the capsid, the attachment protein may be present at varying levels on particles. For example, analyses of SARS-CoV2 and influenza virus particles indicate a heterogeneity in the number of spikes present per virion (6, 7). Further, these types of analyses indicate that these spike proteins are present in varied conformations. It is expected that changes in the number of attachment proteins and their conformation are likely to influence the capacity of the virus to engage host cell receptors and successfully enter cells to launch infection. Yet, the determinants that control this heterogeneity are not fully elucidated.

Mammalian orthoreovirus (reovirus) is a model for studying virus attachment and entry. Reovirus is a nonenveloped virus with a segmented, double stranded RNA (dsRNA) genome (8). Two concentric protein capsids protect the genome of reovirus. The outer capsid, comprised of the μ1, σ3, and σ1 proteins is required for stability in the environment, attachment to host receptors and penetration of the host endosomal membranes. Among these, σ3 and σ1 both serve as attachment proteins (9-13). σ1 forms trimeric fibers that are incorporated into pentameric turrets of λ2 which are found on the icosahedral vertices of the particle (14, 15). Genetic studies link the σ1 encoding S1 segment of reovirus as the determinant of serotype-specific tropism and patterns of disease (16-19). The primary sequences of σ1 proteins of serotype 1 (T1) and serotype 3 (T3) reovirus strains share very little similarity (< 30%)(20). It is therefore thought that engagement of distinct receptors by σ1 proteins of T1 and T3 strains contributes to differences in their tropism.

σ1 is not visualized in CryoEM analysis of reovirus particles (14, 15, 21). Thus, how σ1 is folded or arranged on the particles is not clear. The structures of full-length σ1 molecules released from particles of T1 or T3 reovirus strains or from recombinantly expressed proteins have not been experimentally determined. σ1 forms three distinct structural domains. X-ray crystal structures of individual σ1 domains have been solved. These analyses indicate that despite their sequence divergence, different portions of T1 and T3 σ1 proteins form very similar structures (10, 11, 22, 23). The tail is formed of α-helical coiled coils (22). The body domain is composed of β-spiral repeats (10, 11, 22), and the head domain is comprised of a 8-stranded β-barrel (4, 9). The position of the crystallized domains with respect to each other is not known. While multiple studies suggest that the σ1 fiber is flexible and that this flexibility is important for infection (24, 25), the determinants of this flexibility are poorly defined.

Here, we used ColabFold to model the full-length structures of σ1 from two prototype reovirus strains. Despite their sequence divergence, both σ1 proteins adopt very similar structures. Predictions of structures of chimeric σ1 molecules where the head, body or tail domains are swapped between prototype reovirus strains suggest that chimeric proteins adopt structures that differ in conformation from the parental strains. Confidence metrics associated with the structure predictions suggest that the altered conformations may be related to an expansion in the size of the disordered region that lies between the body and head domain. We present data demonstrating that σ1 molecules with increased disorder or more flexibility compared to the parental strain are poorly incorporated into virions. We also demonstrate that even if the chimeric proteins are efficiently incorporated, their capacity to engage cell surface receptors is altered. Finally, we identify a connection between properties of the μ1 outer capsid protein and the incorporation and conformation of σ1.

## Results

### Predicted structure of full-length σ1

Reovirus T1 and T3 serotypes display differences in tropism and it is thought that this difference relates to the capacity of their σ1 attachment protein to bind distinct host receptors (16-19). For each serotype, the structures of the head domain, head + body domain, and body + tail domain have been solved using X-ray crystallography (4, 9-11, 22, 23). However, structural information about the full-length version of σ1 is not available for any strain. Here, we predicted the full-length structure of type 1 (T1L) and type 3 (T3C44-MA) σ1 proteins using ColabFold where a fast homology search MMSeq2 is paired with AlphaFold2 (26). Because σ1 forms a trimer, we used a trimer for our predictions. As a default, 5 structures were predicted for each construct. For our study here, we focused only on the relaxed structure, in which stereochemical violations and clashes are resolved (27).

The relaxed structures of full-length σ1 trimer from each reovirus serotype appear to be quite similar (Fig. 1C). Further, the previously described head, body and tail domain structures are largely preserved (4, 9-11, 22, 23). The predicted structures are accompanied by a measure of local confidence (27). This confidence is represented by predicted local distance difference test (pLDDT) using scores between 0-100. It is a measure of how well the prediction would agree with an experimentally determined structure. pLDDT analysis of our predicted structures indicate overall high confidence in the predictions (average pLDDT of 83.6 and 81.2 for T1L and T3C44-MA σ1 respectively) (Fig. 1A).

**Fig. 1.**
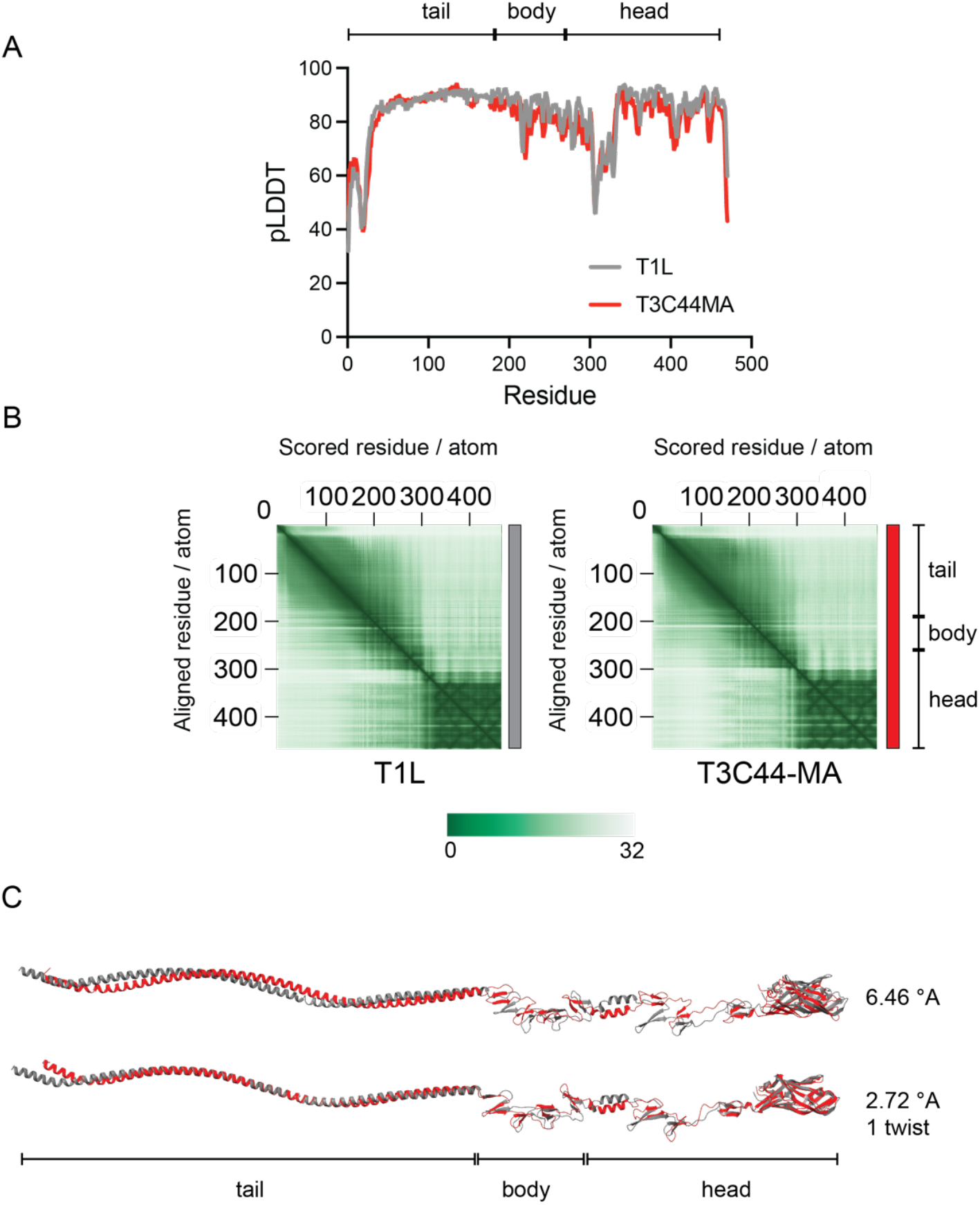
Comparison of ColabFold predicted structures of T1L and T3C44-MA. ColabFold predicted relaxed structures of T1L (grey) and T3C44-MA (red) are compared using (A) pLDDT scores for (B) PAE scores; (C) FATCAT comparison of rigid (top) and flexibly (bottom) aligned structures. RMSD values are indicated. The approximate position of tail, body and head domains of σ1 are shown.

The σ1 protein of T1 strains is 15 amino acids longer than that of T3 strains. Upon alignment of the primary sequence of these proteins it is clear that the T1 strains have an insertion between residues 161 and 174 which makes these proteins longer (28). Though our Colabfold models were generated on the native sequence, comparison of pLDDT values between strains showed that regions of lower pLDDT were offset by about 15 residues (data not shown). To better compare per-residue pLDDT values between T1 and T3 strains, we inserted a gap into the sequence of T3C44-MA between amino acids 161 and 174. This modified version of the sequence from T3C44-MA showed a pLDDT trend similar to that of T1L (Fig. 1A). For σ1 from each strain, we found two regions with low pLDDT scores. First, the N terminus of the protein encompassing region 1-28 has a very low pLDDT. This region of σ1 interacts with the λ2 turret protein (15, 29-31). It is possible that the folding of this region is dependent on σ1-λ2 interaction. Second, the linker region between the body and head domain at residues 305 to 323 has very low pLDDT (pLDDT < 50). For each strain, the number of residues showing lower pLDDT and the relative decrease in pLDDT from the average also are approximately the same.

Accuracy of structure prediction and consequently the associated confidence scores can be affected by the quality of the MSA. MSAs can sometimes include redundant sequences. Thus, simply counting the number of sequences that constitute the MSA doesn’t always provide an accurate measure of their diversity. To address this issue, the concept of “Number of Effective Sequences” (N_eff_) is used, which adjusts for redundancy and better reflects the true diversity or depth of information within an MSA. It is understood that higher N_eff_ results in higher quality prediction (32). To determine whether the low pLDDT values correlated with lower per residue N_eff_, we determined N_eff_ from our MSAs using Neffy (33). For each type of σ1, with the exception of the N terminus of σ1, we find that lower pLDDT did not correlate with lower N_eff_ scores (Fig. S1). Thus, our data suggest that the prominent area of low pLDDT in the head body junction is attributed to the presence of flexible or inherently disordered regions within the σ1 protein. Despite the fact that the primary sequence of T1 and T3 σ1 is divergent, this area of flexibility is conserved. Satisfyingly, this region of low pLDDT within σ1 has also been previously proposed to be flexible (24, 34, 35).

pLDDT is a measure of local confidence. It does not indicate if the residue is in the correct position relative to other parts of the molecule. This information is particularly important for a protein like σ1 that can fold into multiple independent domains. Predicted aligned error (PAE) provides insights into the relative positional error of each residue (27). PAE scores (which range between 0-32) could reveal the position of individual structured domains relative to each other. We find that for both T1L and T3C44-MA σ1, the head domain displays low PAE (Fig. 1B). Similarly, the tail and body domains also display low PAE. However, in each case, the PAE scores of head domain relative to the body and tail domain is high, indicating a lower confidence in their relative positions. The predicted higher PAE between head and body is consistent with the suggestion that this region is flexible. Despite being previously suspected to be modular domains, there appears to be greater confidence in the position of the tail and body domains with respect to each other. These data suggest an absence of flexibility between these domains in parental strains.

Our ability to model the full-length structures of T1L and T3C44-MA σ1 using ColabFold allowed us to compare full-length structures. Traditional structural alignment methods consider proteins to be rigid. Because our analyses above suggested the presence of flexible domains that may allow σ1 to assume different structures, we used <u>F</u>lexible structure <u>A</u>lignmen<u>T</u> by <u>C</u>haining <u>A</u>ligned fragment pairs allowing <u>T</u>wists (FATCAT)(36). In contrast, FATCAT allows rotation and translational movement (twists) within protein so that larger regions can be aligned. Such accommodations are often sufficient to align polymorphic proteins that differ in their conformation. Consequently, FATCAT often yields a lower root mean square deviation (RMSD). We used both rigid and flexible alignment to compare the predicted structures. Since each of the chains that comprise the σ1 trimers are identical, we compared the structure of a single chain. Rigid alignment of T1L and T3C44-MA monomers produced an RMSD value of 6.46 indicating that not all residues are in equivalent positions (Fig. 1C). In contrast, flexible alignment introducing a single twist reduces the RMSD to 2.72. These data suggest that these two proteins assume similar structures but these structures are in a slightly different conformation.

### Chimeric T1L-T3C44-MA σ1 molecules are structurally different from the T1L strain

Previous studies have used chimeric T1 and T3 σ1 proteins to map receptor interaction domains and tropism determinants (37, 38). Such mapping studies could be confounded by changes to the structure of σ1 but how the structures of chimeric σ1 protein differ from those of the parents is unknown. We used ColabFold to predict the structure of chimeric T1L-T3C44-MA σ1 molecules. Overall, we find that each chimeric σ1 is folded into a tail, body, head structure resembling σ1 proteins from the T1 and T3C44-MA parent.

We find that many of the chimeric σ1 proteins display regions of low pLDDT that are larger compared to the parental viruses (Fig. 2). For 1-1-3 (T1 tail-T1 body-T3 head), we find that pLDDT decreases at residue 263 (instead of 305 for the parental strains) and remains low until about 316. The start of the decrease appears to coincide with the junction between T1 body and T3 head. There is also an additional dip in pLDDT values at residues 396-402 within the head region. For 1-3-1, the pLDDT values decrease between residues 282-303. While this region of lower confidence is smaller than that predicted for 1-1-3, like the 1-1-3 chimera, in this case, the reduction in pLDDT also occurs in a region that is the junction between T1 body and T3 head. For 3-1-1 σ1, the values are very similar to parental strain T1L with no pronounced extension of the region with low pLDDT. For 3-1-3 σ1, a larger portion of σ1 is predicted to have lower pLDDT (residues 236-304). Interestingly, like the examples above a reduction in pLDDT is observed starting at the body and head junction where in this case a T1 body is appended to a T3 head. We also observe a decrease in pLDDT within the head region. For 3-3-1, lower pLDDT values are predicted between residues 246-295, once again falling starting in the region where the T1 body is appended to the T3 head. Finally for 1-3-3, we find a somewhat unusual pLDDT score pattern. Unlike parental T1L, we find a very small region of pLDDT decrease between the body and head junction. We see an additional sharp decrease in pLDDT at residue 400, which falls within the head region. Because pLDDTs correlate with MSA quality, we evaluated whether decreased in pLDDT was due to lower N_eff_ score. While the lower pLDDT near the N-terminus of σ1 and the sharp decrease in pLDDT within the head region of some chimeras correlated with lower N_eff_, changes in the head-body junction of chimeric σ1 did not (Fig. S1). pLDDT values obtained from these predictions therefore suggest that that a mismatch between head and body domain results in a more flexible structure. Interestingly, a mismatch between the tail and body domain does not have a significant effect on pLDDT values.

**Fig. 2.**
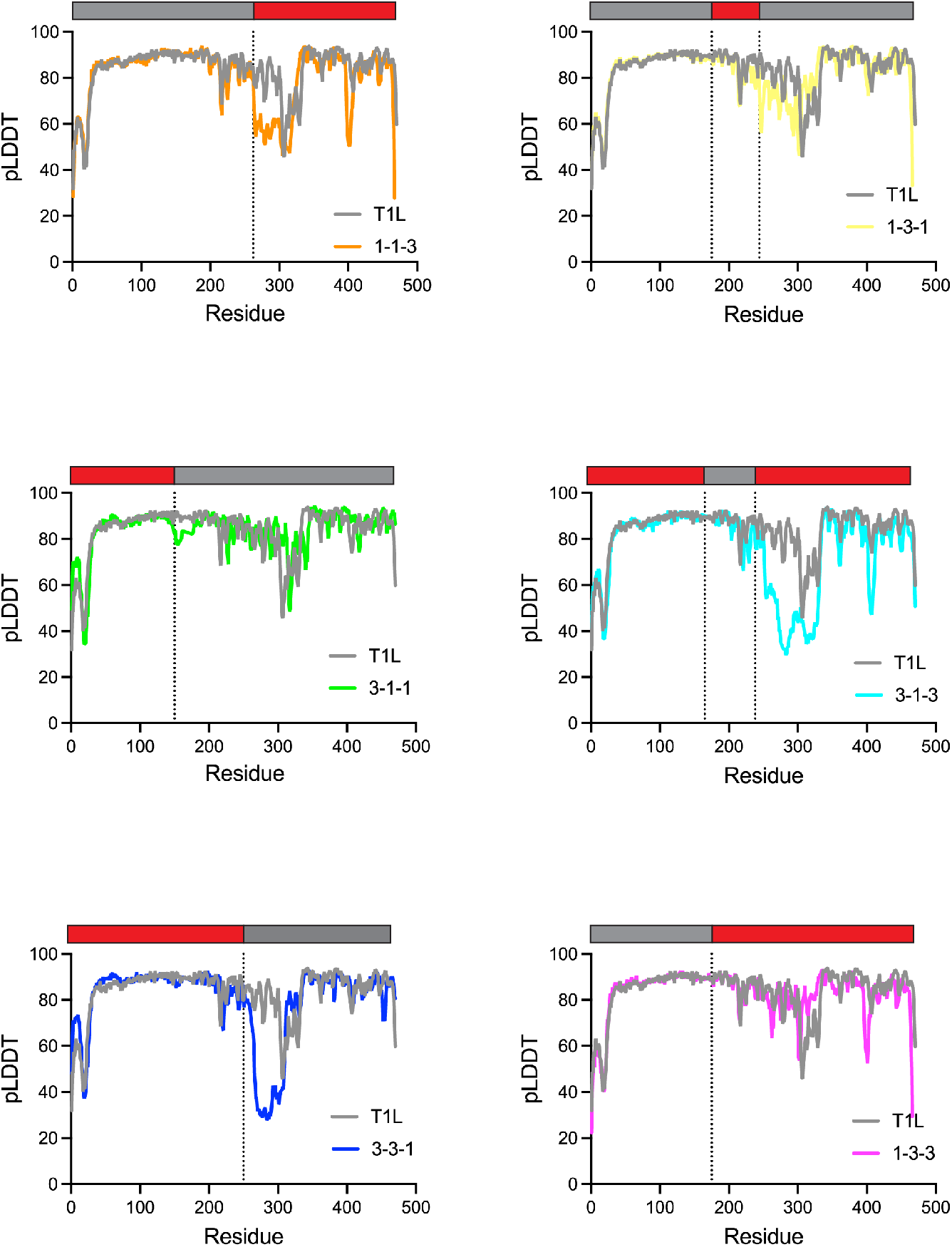
Comparison of pLDDT scores of chimeric σ1. pLDDT scores of chimeric σ1 are compared to T1L. Schematics of chimeras are shown above with T1 and T3 derived portions in grey and red respectively. Dotted lines indicate junctions between T1L and T3C44-MA domains.

PAE analysis of chimeric σ1 also suggest structural differences between most chimeric σ1 protein and parental T1L σ1. Similar to T1L σ1, though the head domain displays low PAE, its position relative to the body domain is not confidently predicted for any chimera (Fig. 3).

**Fig. 3.**
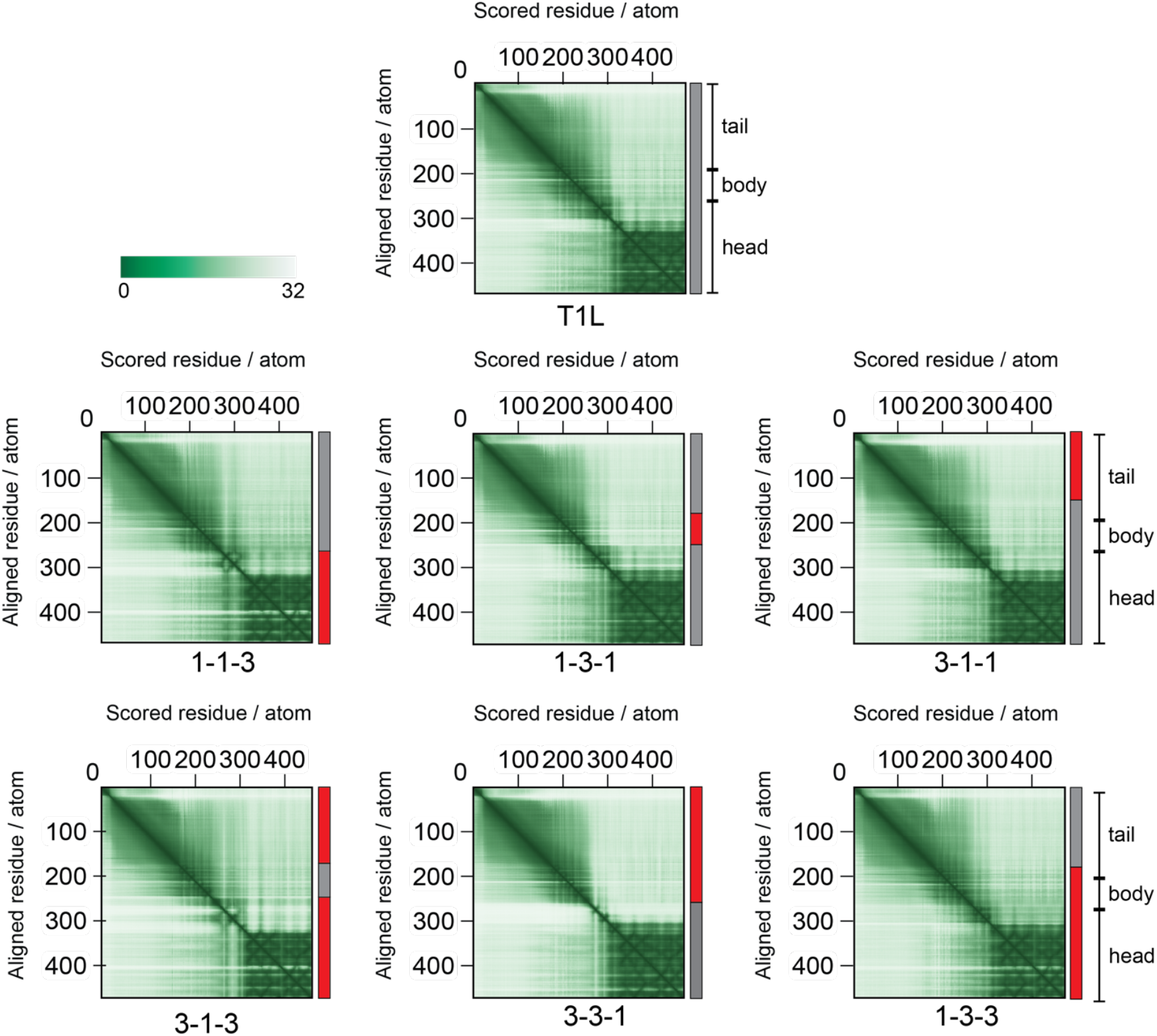
Comparison of PAE scores of chimeric σ1. PAE plots of T1L and chimeric σ1 are shown. Schematics of chimeras are shown above with T1 and T3 derived portions in grey and red respectively. The approximate position of tail, body and head domains of σ1 are also shown.

Further examination of PAE plots of chimeric σ1 indicate that with the exception of 3-1-1 and 1-3-3, we observe a more distinct separation between the tail and body domain. Thus, for 1-1-3, 1-3-1, 3-1-3, 3-3-1, the position of the body domain relative to the tail domain also may be different. Together, these results indicate that many of the chimeras vary in structure of σ1 compared to the parental virus.

We also compared the structures of the chimeric σ1 molecules to those of the parent using FATCAT using both rigid and flexible alignment (Fig. S2 for rigid alignment and Fig. 4). With the exception of 1-3-1 σ1, which aligned with equivalent RMSD under both conditions, each chimera showed a higher RMSD when aligned as a rigid body. Each of the structures that aligned better when flexible alignment required one or more twists to attain better alignment.

**Fig. 4.**
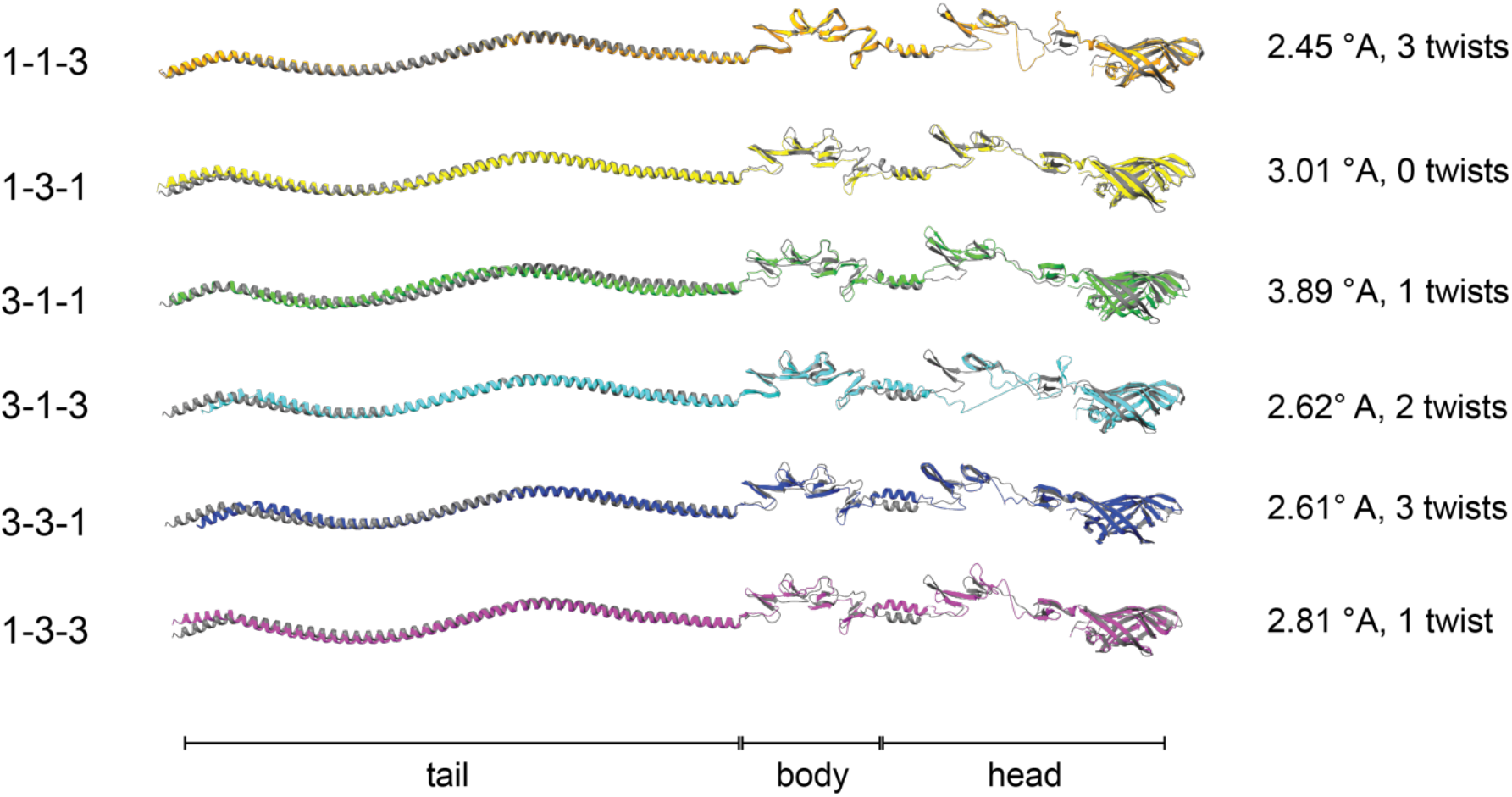
Structural alignment of chimeric σ1. T1L σ1 (grey) is aligned with chimeric σ1 molecules using flexible alignments. RMSD values and the number of twists needed for better alignment of flexible structures are shown. The approximate position of tail, body and head domains of σ1 are also shown.

These analyses again suggest that chimeric σ1 molecules assume structures that vary in conformation.

### Incorporation efficiency of chimeric σ1 molecules differs from the parental strain

Our results presented above based on ColabFold predictions suggest differences in the structures of chimeric σ1 proteins. While it is not known exactly what type of structural features in σ1 are necessary for its incorporation into virions, it is possible that σ1 structure might influence its incorporation. To address this possibility, we used previously generated recombinant T1L viruses that differ from each other only in their σ1 sequence (38). Interestingly, a previous study on these viruses reported that a T1L strain expressing 1-3-3 could not be recovered. To evaluate σ1 encapsidation on the virion, equal amount of purified particles of each strain were resolved on denaturing SDS-PAGE gel. All recombinant strains show equal stoichiometry of the major viral capsid proteins (λ, μ1, σ2, σ3). Given the differences in primary sequence of σ1 for T1 and T3 viruses, no σ1-specific antibodies can be used to quantify σ1 levels in all virus strains. We therefore quantified σ1 levels in the preparation by measuring the intensity of Commassie Blue stain signal. Compared to T1L, each chimeric virus appeared to contain slightly lower level (30-70%) of σ1 (Fig. 5A).

**Fig. 5.**
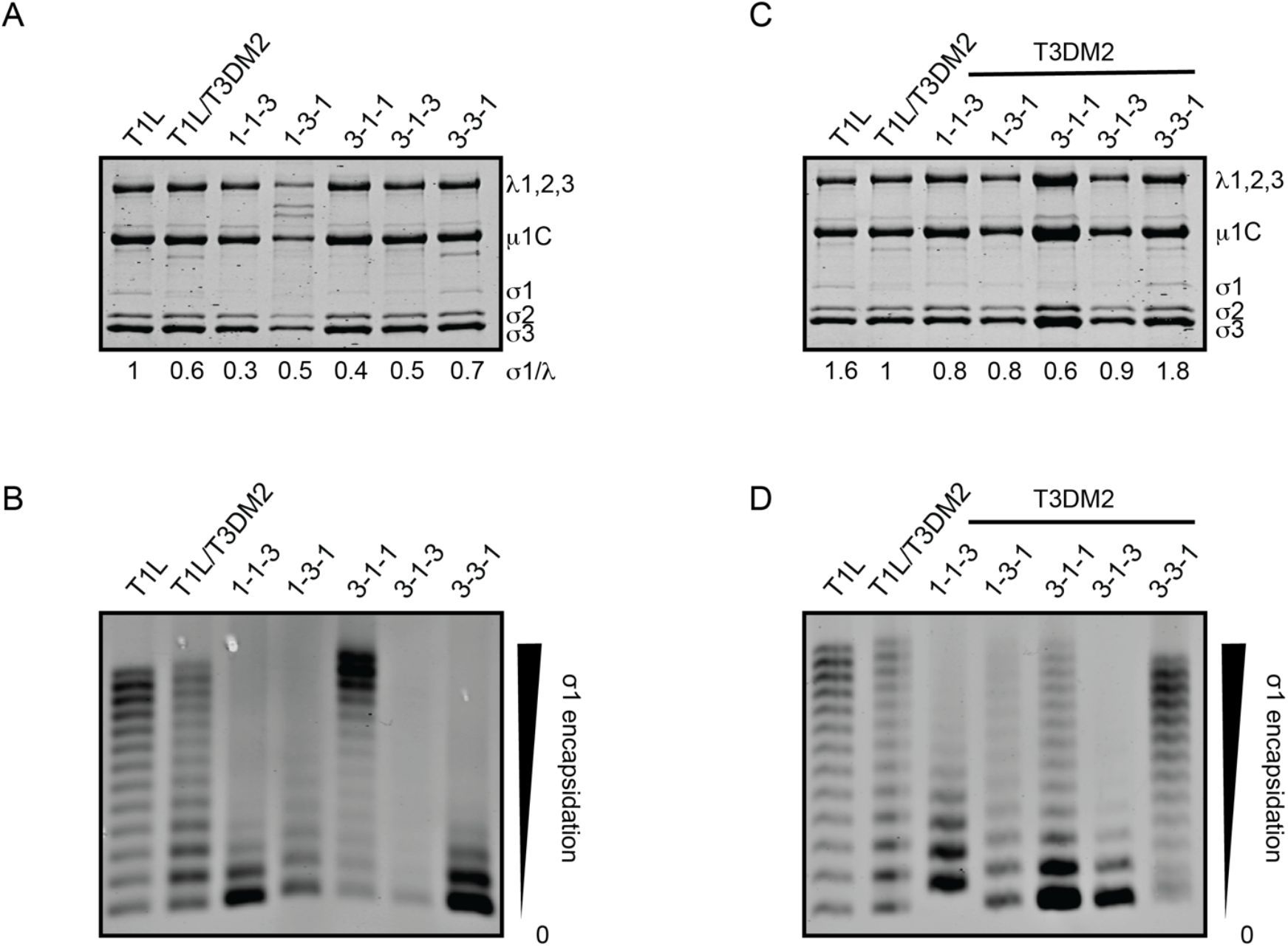
Encapsidation of σ1 is altered in viruses expressing chimeric σ1. (A,C) 1×10^11^ particles of the indicated virus strains were resolved on SDS-PAGE gels and stained with Coomassie brilliant blue. For each virus, band intensities of the region encompassing the σ1 band were divided by the intensity of the λ band. Intensity of the parental virus is indicated is set to 1. (B,D) 1×10^11^ particles of the indicated virus strains were resolved on 2% agarose and stained with colloidal blue stain. For each virus, the intensity of each band was compared to the total intensity of the entire lane. The encapsidation patterns are representative of encapsidation from at least 2 independent purified virus preparations.

While the above results measure the total amount of σ1 on the particles, populations of purified reovirus particles are heterogenous. Whereas some particles contain no σ1 molecules incorporated onto the particle, others can contain up to 12 σ1 trimers (39). Some strains contain a large population of particles with high level of incorporated σ1, others, especially mutant viruses, can have a large population of σ1-less particles (30, 31, 40, 41). To evaluate whether chimeric σ1 molecules are incorporated at the same efficiency as T1L σ1, we resolved particles on native agarose gels, which separate particles based on the level of σ1 incorporation. For T1L, we find that the population contains particles with the entire range (0-12) of incorporated σ1 trimers (Fig. 5B). In comparison to T1L, viruses with 1-1-3, 1-3-1, and 3-3-1 appear to have more particles with lower levels of σ1. In contrast, 3-1-3 (which appears as a faint smear) and 3-1-1 contains particles that span an entire range of σ1 incorporated. The population of 3-1-1 in particular contains particles with very high level of incorporated σ1. These data suggest that change to any of the domains of σ1, including the body and head, which are not known to make interactions with the remainder of the particle, alter incorporation efficiency. In context of our structure predictions, it is possible that these differences in incorporation are a consequence of altered conformation of σ1 which prevents it from becoming incorporated or remain stably assembled on the particle.

Previously, we demonstrated that properties of μ1 impact the function of σ1, likely due to effects on its conformation (42). To test our idea that conformation of chimeric σ1 may impact its virion incorporation efficiency, we generated recombinant viruses with 8 genes from T1L, the μ1-encoding M2 gene from T3D and each of the chimeric σ1 constructs (except 1-3-3). In this genetic background, the stoichiometry of the major viral proteins was preserved. We compared the level of σ1 incorporation for chimeric σ1 proteins to σ1 in T1L/T3DM2. We find that a majority of the chimeric σ1 molecules are incorporated with efficiency similar to this parent.

Incorporation of the 3-3-1 σ1 chimera was modestly lower than that for T1L/T3DM2 (Fig. 5C). To further characterize this effect, we evaluated σ1 incorporation onto the particles using native agarose gel electrophoresis. Compared to T1L, the σ1 encapsidation pattern of T1L/T3DM2 is different (Fig. 5D). While we observe that the preparation contains virions with an entire range of σ1 incorporation, there appear to be fewer particles that incorporate an intermediate level of σ1 and more particles that incorporate lower levels of σ1. We also compared incorporation of σ1 chimeras to σ1 on T1L/T3DM2. We observe that each chimera differs from T1L/T3DM2. For 1-1-3, 1-3-1, 3-1-1, and 3-1-3, the population distribution is shifted towards particles with lower σ1. In contrast, for 3-3-1, the population contains virions with a higher level of σ1. Like the results for T1L, these data also indicate that changes to σ1 sequence alter its incorporation levels on virions. Additionally, when encapsidation of σ1 chimeras are compared between T1L and T1L/T3DM2 genetic backgrounds, we find that encapsidation patterns of 3-1-1, 3-1-3, and 3-3-1 σ1 are different depending on the background of the M2 gene. In contrast 1-1-3 and 1-3-1 show similar patterns in both T1L and T1L/T3DM2 background. These data indicate that the encapsidation of the T3 virus derived tail domain containing σ1 is affected by the nature of the μ1 protein.

### Viruses with chimeric σ1 display differences in attachment to cells

The σ1 protein plays a critical role in viral attachment to the cell. Though the relationship between σ1 incorporation levels and infection efficiency is not linear (39), levels of encapsidation have the potential to affect viral attachment to cells. Even under conditions where encapsidation levels are not different, attachment could be affected if the conformation of encapsidated σ1 differs. We therefore tested with viruses bearing chimeric σ1 attachment proteins showed difference in attachment using a plate based attachment assay. Compared to T1L, each virus with the exception of 3-3-1, displayed a lower efficiency of attachment (Fig. 6A). Compared to the encapsidation data shown above (Fig. 5A), these attachment data indicate that there appears to be no correlation between the σ1 encapsidation pattern and attachment efficiency. For example, 3-3-1 and 1-1-3 have similar encapsidation pattern despite displaying different attachment efficiency. Similarly, we find that despite containing a population of particles with high levels of σ1 encapsidation, 3-1-1 has a lower attachment efficiency.

**Fig. 6.**
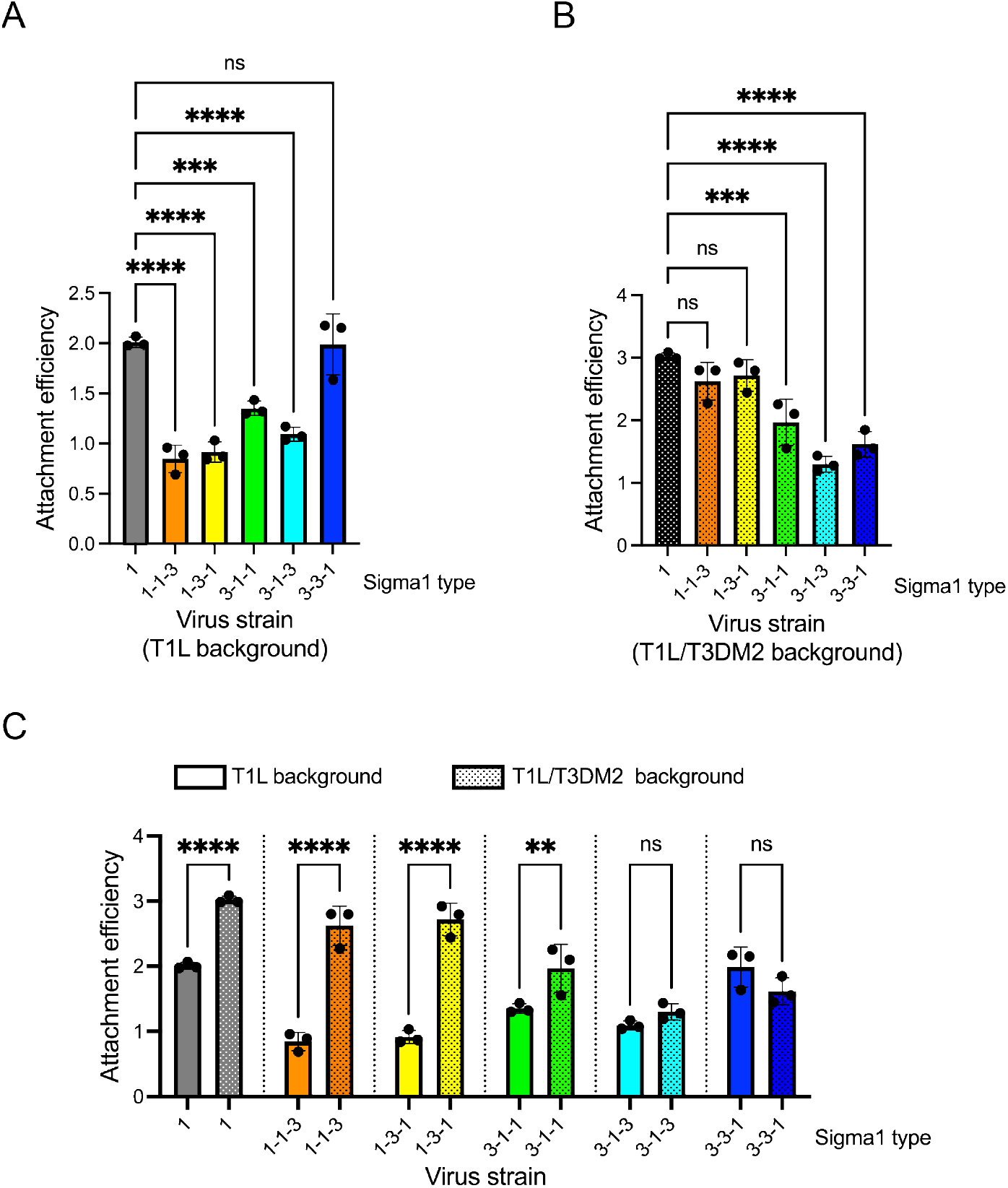
σ1 chimeras display differences in attachment efficiency. L929 cells were adsorbed with 5 x 10^4^ virus particles per cell of the indicated viral strains. Attached virus was detected using reovirus-specific rabbit polyclonal antisera and secondary antibody. Cells were counter stained with a DNA stain. Viral attachment was quantified using infrared scanner. Results are expressed as mean binding index (ratio of fluorescence of attached virus to that of cellular DNA) for three independent samples. Error bars indicate SD. *, P < 0.05 as determined by ANOVA with Bonferroni multiple comparison test in comparison to T1L (A) or T1L/T3DM2 (B). (C) Attachment efficiency of viruses in T1L and T1L/T3DM2 background is compared. *, P < 0.05 as determined by ANOVA with Šídák’s multiple comparison test in comparison to the virus containing the same S1 but different M2 gene.

We also measured attachment efficiency of σ1 chimeras generated in the T1L/T3DM2 background. We find that in comparison to T1L/T3DM2, the attachment of 1-1-3 and 1-3-1 were unaffected (Fig. 6B). In contrast, the attachment of 3-1-1, 3-1-3 and 3-3-1 was lower. Comparison of these attachment data with σ1 encapsidation profile of these viruses indicate once again that encapsidation levels do not match attachment efficiency. Whereas 1-1-3, 1-3-1, 3-1-1, and 3-1-3 all contain more particles with lower level of σ1, only some of them (3-1-1 and 3-1-3) have weaker attachment. Similarly, despite having more particles with high σ1 encapsidation, 3-3-1 fails to attach efficiently.

In previous work, we demonstrated that T1L/T3DM2 attached to cells more efficiently than T1L (43). However, T1L/T3D S1M2 did not attach more efficiently than T1L/T3DS1. These findings suggested that the T3D μ1 mediated enhancement of attachment requires the presence of the T1L σ1 protein. We used our chimeric viruses to map regions in σ1 that are important for T3D μ1 mediated effect on attachment. Because the attachment experiments shown above in Fig. 6A and 6B were performed concurrently, we were able to compare the attachment efficiency of all viruses. In Fig. 6C, the same data shown above is represented differently. Consistent with our previous observation, T1L/T3DM2 shows greater attachment than T1L. A similar trend was also observed for 1-1-3 and 1-3-1. In contrast, the attachment increase was less convincing (for 3-1-1) or absent (for 3-1-3 and 3-3-1). Based on these data, we conclude that the presence of the T1 tail is necessary for T3D μ1 mediated enhancement of attachment.

## Discussion

Despite its importance as a viral attachment factor and its role in controlling the tropism of reovirus, structural information on the full-length structure of σ1 is lacking. Here, we use ColabFold to predict the structures of full-length σ1 from type 1 and type 3 strains. This analysis reveals a region of flexibility between the body and head domain of σ1 for each serotype (Fig. 1). ColabFold analysis of chimeric σ1 molecules comprised of T1 and T3 σ1 domains indicate that these proteins assume a structure that displays some differences from the structure of the parental σ1 (Fig. 2, 3 and 4). In particular, chimeric σ1 display more disorder suggesting more flexibility in the head-body junction if these regions are mismatched. We demonstrate that change in flexibility of σ1 affects its encapsidation on virions (Fig. 5). We also demonstrate that independent from attachment, the differences in structures of chimeric σ1 results in altered cell attachment efficiency of these viruses (Fig. 6). Finally, we demonstrate that the μ1 protein controls both encapsidation efficiency and attachment properties of σ1 (Fig. 5 and 6). These findings are summarized in Table 1.

**Table 1:** Summary of characterization of virus strains. Predicted area of flexibility is derived from analyses shown in Figure 2. Encapsidation results are from experiments in Figure 5. Attachment data are derived from Figure 6.

| Virus strain | Predicted area of flexibility | $\sigma 1$ Encapsidation | | Virion Attachment | |
| --- | --- | --- | --- | --- | --- |
| | | T1L $\mu 1$ | T3D $\mu 1$ | T1L $\mu 1$ | T3D $\mu 1$ |
| T1L | ++ | ++++ | ++++ | ++++ | ++++ |
| 1-1-3 | ++++ | ++ | ++ | ++ | ++++ |
| 1-3-1 | +++ | ++ | ++ | ++ | ++++ |
| 3-1-1 | ++ | +++++ | ++ | +++ | +++ |
| 3-1-3 | ++++ | ++ | ++ | +++ | ++ |
| 3-3-1 | ++++ | ++ | ++++ | ++++ | ++ |
| 1-3-3 | + | ND | ND | ND | ND |

The σ1 protein is divided into three structural domains - tail, body and head. While recombinant versions of individual σ1 domains and some adjacent σ1 domains have been subjected to structural analysis by crystallography (4, 9-11, 22, 23), the structure of full-length σ1 remains unknown. Further, since crystal structures obtained represent one possible conformation of σ1, it is unclear if this is the only conformation σ1 can assume. In negative stained electron micrographs, σ1 fibers tipped with a knob protrude from the particle (44). Not all σ1 molecules on a particle appear to be of the same length suggesting structural flexibility. Consistent with this, σ1 molecules released from viral particles can adopt alternate structures with kinks between the knob and fiber and also along the length of the fiber (34). Based on computational processing of electron micrographs and sequence analysis, it was predicted that flexible interdomain regions exist between the tail and body and between the body and head (34). Molecular dynamics simulations of σ1 also suggest mobility of the head domain with respect to other parts of the molecule (35). Our ColabFold predictions of full-length type 1 and type 3 σ1 protein and the associated pLDDT scores suggest that the region between the body and head is flexible (Fig. 1). Our analysis did not predict a flexible region between the tail and body domain. This idea is consistent with more recent structural evidence which indicates that the tail seamlessly transitions to the body (22). For parental type 1 and type 3 strains the region of flexibility between the body and head domain is structurally conserved. pLDDT analyses of chimeric σ1 constructs indicate that amino acids sequences on both sides of the junction regulate how large the flexible portion is and the extent to which it is flexible (Fig. 2 and 3). Our findings indicate that when these regions are mismatched between strains, the flexibility of the molecule increases. Thus, our work reveals determinants that control the extent of this flexibility.

The assembly process of reovirus is partially understood. In infected cells, progeny viral cores are found in viral factories (45, 46). These factories are spatially separated from newly synthesized outer capsid proteins (which includes σ1 along with σ3 and μ1) because those proteins associate with lipid droplets. These compartment later merge to generate virions. In *in vitro* experiments, outer capsid proteins expressed in insect cell lysate can be recoated onto purified cores to generate infectious particles (47). σ1 encapsidation into λ2 turrets requires their closure from their open conformation present in cores (14). This closure is expected to be mediated by the presence of σ3-μ1 heterohexamers on the particle. However, the order of events during outer capsid assembly has not been investigated. Electron microscopic evidence indicate that each particle has multiple σ1 molecules arranged with regular spacing between them but the number of σ1 proteins on each particle varies (44). This finding is corroborated by evidence that in a given purified virus preparation, particles bearing different levels of σ1 are present (39). Single amino acid substitutions in the N-terminal portion of the tail and in the head region alter its encapsidation pattern (30, 31, 40). It has been experimentally demonstrated that the N-terminus of the σ1 tail is important for encapsidation because it interacts with the λ2 protein (29). However, it is not clear how changes in the head region can influence encapsidation. One possible explanation for impact of this distal region on encapsidation is that point mutations in σ1 alter its conformation. Indeed, one substitution that alters incorporation into virions is thought to assume an altered conformation (41, 48). This idea is supported by evidence of reduced susceptibility of the protein to a protease even though the sequence of the cleavage site is intact. Experimental information indicating the extent to which this change alters structure is lacking. Given that structure prediction algorithms are not sensitive to point mutations, this information cannot be obtained computationally. Our data presented here using ColabFold structure prediction of chimeric σ1 also corroborate the idea that σ1 structure regulates its encapsidation. Our results demonstrate the importance of σ1 flexibility at the body-head junction. We find that when flexibility is similar to that of parental T1L, particles with highly encapsidated σ1 are assembled (eg. 3-1-1) (Fig. 2 and 5, Table 1). In contrast when the area of flexibility is expanded, we find more particles with lower σ1 encapsidation (eg, 1-1-3,1-3-1, 3-1-3, and 3-3-1). Finally, when σ1 flexibility in the head-body junction is decreased (1-3-3), infectious viruses cannot be recovered suggesting a potential encapsidation defect.

In this study, we demonstrate that properties of μ1 influence the encapsidation of σ1 molecules. On assembled virus particles, μ1 and σ1 do not contact each other directly (14, 15, 21). μ1 subunits arranged around the particle vertex interact with λ2, which forms turrets into which σ1 is inserted. Thus, when properties of μ1 change, its interaction with λ2 could be qualitatively different. Such a change could subtly change the structure of λ2 and thereby affect whether the σ1 protein can become encapsidated or remain encapsidated. Similarly, changes to the structure of λ2 could also affect its interaction with the encapsidated σ1 and alter how it is presented on virions. Because our predictions suggest that chimeric σ1 molecules fold differently, it is possible that some of these altered structures are accommodated more easily than others. Such a possibility could produce μ1 dependent effects on the encapsidation of some but not all chimeric σ1. We also demonstrate that independent of encapsidation, properties of μ1 also influence cell attachment function of σ1. We surmise that μ1 dependent changes in λ2 structure could impact its interaction with σ1 via the tail region and alter how σ1 is presented on the particle. Our attachment data presented here indicate that the presence of the type 1 σ1 tail is sufficient for T3 μ1 mediated enhancement of the attachment function of σ1.

This result lends more support for the idea that μ1 could indirectly affect the interaction of σ1 with λ2.

## Materials and Methods

### ColabFold, N_eff_ and FATCAT analysis

Prediction of structure of σ1 trimers was done using local copy of ColabFold installed on Big Red 200, an HPE Cray EX supercomputer (26). The following parameters were used: “num_queries”:1,”use_templates”:false,”num_relax”:1,”relax_max_iterations”:2000,”relax_tolera nce”:2.39,”relax_stiffness”:10,”relax_max_outer_iterations”:3,”msa_mode”:”mmseqs2_uniref_en v”,”model_type”:”alphafold2_multimer_v3”,”num_models”:5,”num_recycles”:30,”recycle_early_s top_tolerance”:0.4,”num_ensemble”:1,”model_order”:[1,2,3,4,5],”keep_existing_results”:true,”ra nk_by”:”multimer”,”max_seq”:508,”max_extra_seq”:2048,”pair_mode”:”unpaired_paired”,”pairin g_strategy”:”greedy”,”host_url”:”https://api.colabfold.com“,”user_agent”:”colabfold/1.5.5(fdf3b23 5b88746681c46ea12bcded76ecf8e1f76)”,”stop_at_score”:100,”random_seed”:0,”num_seeds”:1 ,”recompile_padding”:10,”commit”:”fdf3b235b88746681c46ea12bcded76ecf8e1f76”,”use_dropo ut”:false,”use_cluster_profile”:true,”use_fuse”:true,”use_bfloat16”:true,”version”:”1.5.5”

MSAs data (.a3m files) obtained from ColabFold were analyzed were analyzed using Neffy to obtain per residue N_eff_ scores (33).

Confidence scores were measured in ColabFold using pLDDT (predicted local distance difference test) and PAE. Structures were visualized using ChimeraX (49). Pairwise comparison of structures was completed using FATCAT using both rigid and flexible alignment(36).

### Cells

Spinner adapted Murine L929 cells were maintained in Joklik’s MEM (Lonza) supplemented to contain 5% fetal bovine serum (FBS) (Sigma-Aldrich), 2 mM L-glutamine (Invitrogen), 100 U/ml penicillin (Invitrogen), 100 μg/ml streptomycin (Invitrogen), and 25 ng/ml amphotericin B (Sigma-Aldrich). Spinner adapted L929 cells were used for cultivating, purifying and titering viruses. L929 cells obtained from ATCC were maintained in Eagle’s MEM (Lonza) supplemented to contain 5% fetal bovine serum (FBS) (Sigma-Aldrich), 2 mM L-glutamine (Invitrogen). Experiments to measure attachment, infectivity, and cell death were done using L929 cells form ATCC.

### Generation of Recombinant Viruses

Strain T1L was re-derived from plasmids using reverse genetics. σ1 chimeric viruses in the T1L background were previously described and were kindly shared by Terry Dermody (University of Pittsburgh)(38). Viruses in the T1L/T3DM2 background were generated by plasmid based reverse genetics using chimeric S1 constructs obtained from the Dermody laboratory. To confirm sequences of mutant viruses, viral RNA was extracted from infected cells and subjected to reverse transcription PCR using multiple sets of S1-specific primers. PCR products were resolved on Tris-acetate-EDTA agarose gels, purified and confirmed by sanger sequencing.

### Purification of viruses

Purified reovirus virions were generated using second- or third-passage L-cell lysates stocks of reovirus. Viral particles were Vertrel-XF (Dupont) extracted from infected cell lysates, layered onto 1.2-to 1.4-g/cm^3^ CsCl gradients, and centrifuged at 187,183 x *g* for 4 h. Bands corresponding to virions (1.36 g/cm^3^) were collected and dialyzed in virion-storage buffer (150 mM NaCl, 15 mM MgCl^2^, 10 mM Tris-HCl [pH 7.4] (50). The concentration of reovirus virions in purified preparations was determined from an equivalence of one OD unit at 260 nm equals 2.1 x 10^12^ virions/ml (51).

### Agarose gel separation of reovirus particles by σ1 content

1 x 10^11^ virus particles were resuspended in dialysis buffer, mixed with 2X Gel loading dye (NEB) and resolved on 1% ultra-pure agarose gel (Invitrogen) in 1X TAE pH 7.2 at constant 25 V for 18 h. The gel was stained with NOVEX colloidal blue staining kit (Invitrogen) for 6 h and destained overnight in water. Gel was scanned using Odyssey Infrared Imager (LI-COR).

### Assessment of reovirus attachment

L929 cells (4 x 10^4^ cell per well) grown in 96-well plates were chilled at 4°C for 15 min and then adsorbed with particles of the indicated virus strains at 4°C for 1 h. Cells were washed with chilled PBS and blocked with PBS-BSA at 4°C for 15 min. Cells were then incubated with polyclonal rabbit anti-reovirus serum at a 1:2500 dilution in PBS-BSA at 4°C for 30 min. The cells were washed twice with PBS-BSA followed by incubation with 1:1000 dilution of Alexa Fluor 750 labeled goat anti-rabbit antibody at 4°C for 30 min. After two washes with PBS-BSA, cells were stained with 1:1000 dilution of DNA stain, DRAQ5 (Cell Signaling technology), at 4°C for 5 min. Cells were washed and then fixed with 4% formaldehyde at room temperature for 20 min. Fluorescence intensity was measured using the Odyssey Imaging System and the Image Studio Lite software (LI-COR). For each well, the ratio of fluorescence at 800 nm (for attached reovirus) and 700 nm (for total cells) was quantified. Attachment efficiency in arbitrary units was quantified using the following formula. Attachment efficiency = (Green/Red)_infected_ - (Green/Red)_uninfected_.

---

**Fig. S1.**
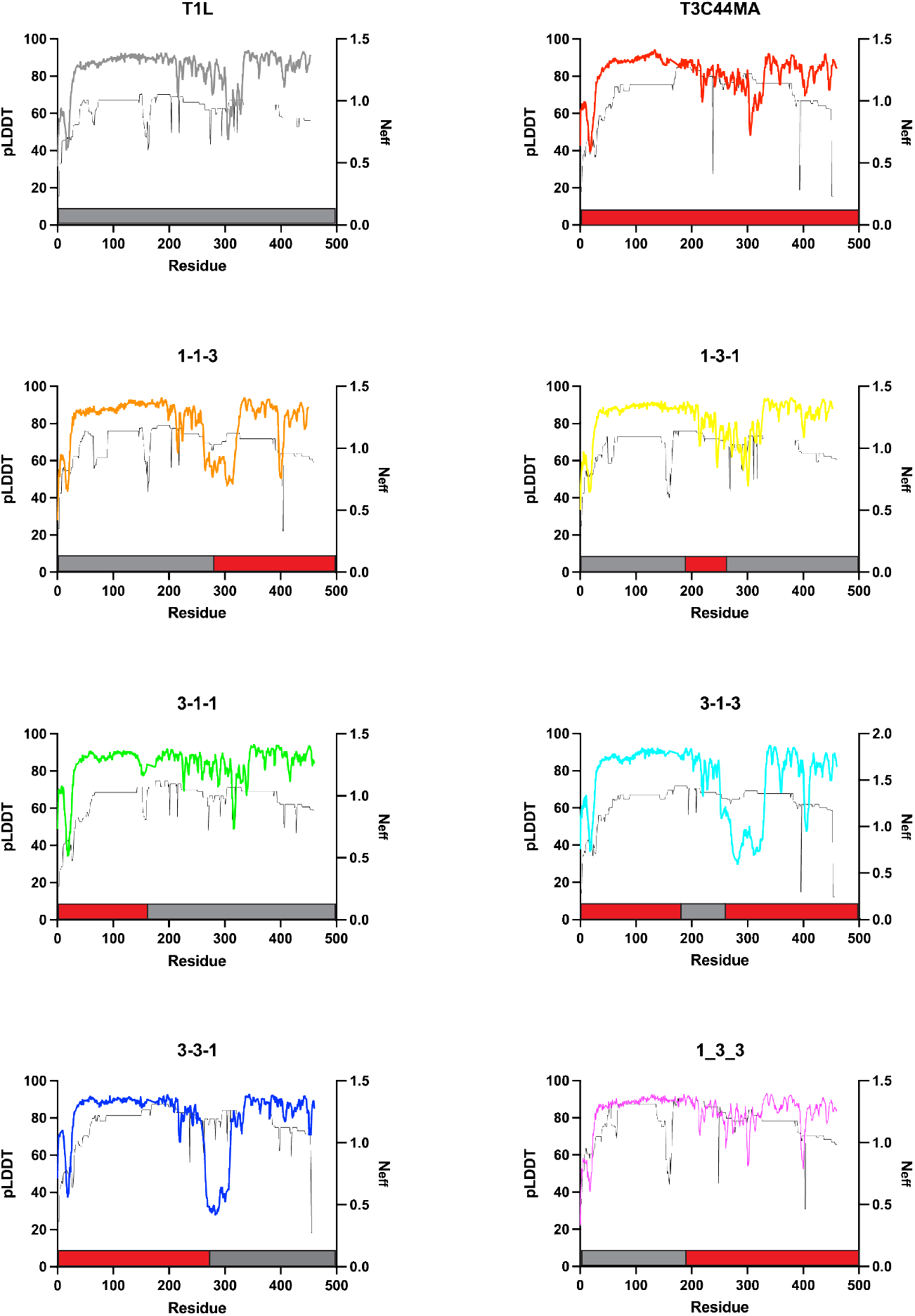
Relationship between pLDDT and N_eff_ scores. pLDDT (thick colored lines) and N_eff_ scores (thin black line) of the indicated σ1 protein were compared. Schematics of chimeras are shown above with T1 and T3 derived portions in grey and red respectively.

**Fig. S2.**
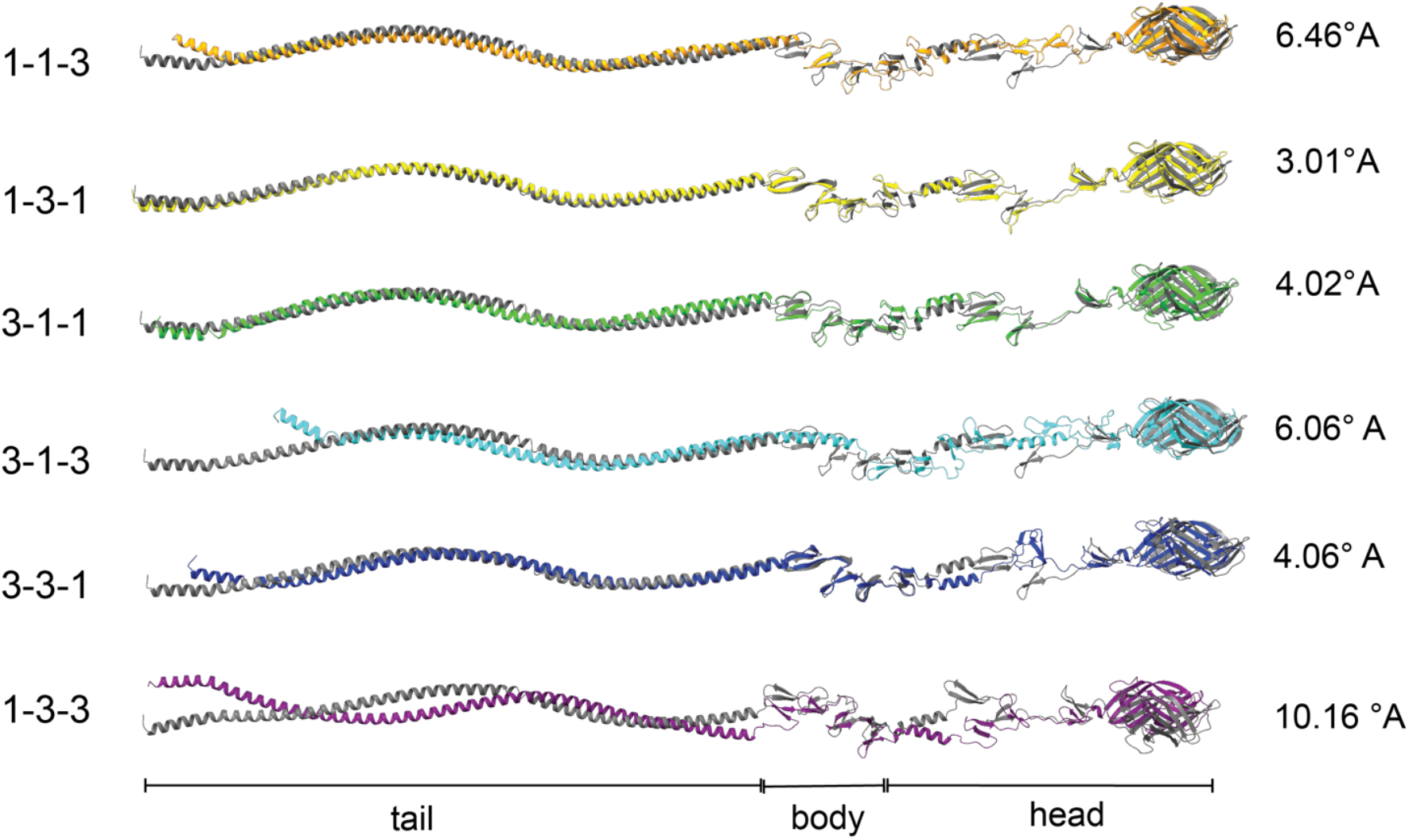
Rigid structural alignment of chimeric σ1. T1L σ1 (grey) is aligned with chimeric σ1 molecules using rigid alignments. RMSD values are shown. The approximate position of tail, body and head domains of σ1 are also shown.

